# QC-GN^2^oMS^2^: a Graph Neural Net for High Resolution Mass Spectra Prediction

**DOI:** 10.1101/2023.01.16.524269

**Authors:** Richard Overstreet, Ethan King, Julia Nguyen, Danielle Ciesielski

## Abstract

Predicting the mass spectrum of a molecular ion is often accomplished via three generalized approaches: rules-based methods for bond breaking, deep learning, or quantum chemical (QC) modeling. Rules-based approaches are often limited by the conditions for different chemical subspaces and perform poorly under chemical regimes with few defined rules. Quantum chemical modeling is theoretically robust but requires significant amounts of computational time to produce a spectrum for a given target. Among deep learning techniques, graph neural networks (GNNs) have performed better than previous work with fingerprint-based neural networks in mass spectral prediction.^1^ To explore this technique further, we investigate the effects of including quantum chemically derived features as edge features in the GNN to increase predictive accuracy. The models we investigated include categorical bond order, bond force constants derived from Extended Tight-Binding (xTB) quantum chemistry, and acyclic bond dissociation energies. We evaluated these models against a control GNN with no edge features in the input graphs. Bond dissociation enthalpies yielded the best improvement with a cosine similarity score of 0.462 relative to the baseline model (0.437). In this work we also apply dynamic graph attention which improves performance on benchmark problems and supports the inclusion of edge features. Between implementations, we investigate the nature of the molecular embedding for spectral prediction and discuss the recognition of fragment topographies in distinct chemistries for further development in tandem mass spectrometry prediction.

## Introduction

Liquid chromatography tandem mass spectrometry (LC-MS/MS) techniques are widely used in chemical identification for non-target analysis of environmental pollutants, proteomics, metabolomics, and drug characterization.^2–10^ Mass spectrometry instruments designed for these applications commonly use electrospray ionization (ESI) to introduce an analyte for characterization. Post-ionization target molecular ion or precursor peaks are fragmented via collision induced dissociation (CID) at low (0-100 eV) or high (keV) energies.^11^ High-resolution detection of molecular fragments is often accomplished via time-of-flight (ToF) measurements or Orbitrap current analysis. ^12^ When correctly applied to a chemical system, these techniques are invaluable for characterization.

Despite the success of LC-MS/MS techniques, identification of unknown compounds and spectral prediction remain challenging problems. One of the most practical methods for compound identification remains comparing experimental spectra to spectra available in existing reference libraries including NIST, MassBank, Metlin, and MoNA.^13–16^ Matching unknowns to identified spectra in reference libraries can provide insight into chemical structure. However, due to the cost of chemical standards and the wide range of instrument parameters/configurations, spectral databases are unable to encompass a chemical/parameter space large enough for the identification of all novel compounds. To address this pitfall, data informed projects such as CFM-ID, CSI-FingerID, and MAGMA^17–19^ aim to fill this void in database comparison capability. Projects such as Mass Frontier use rules-based methods for chemical fragmentation. These rule sets are informed via prior experience in the literature or sourced from in-house databases where instrument and sample conditions can be carefully controlled. While useful for compounds that fall within the purview of existing rules, these methods fail for novel chemistries. Other more exhaustive approaches include tools such as CFM-ID, which aim to combinatorially model fragmentation to predict a MS/MS spectrum.^17^

Computation of mass spectra from first principles was originally implemented in the work of Grimme et al. for electron impact mass spectrometry (EI-MS).^20^ Under this approach, target molecules are modeled using molecular dynamics and forces are computed by an approximate quantum mechanics method. This allows for explicit modeling of fragmentation pathways and aggregation into an *in silico* spectrum without dependence on reference library information. The approach of modeling from first principles has been implemented in the QCxMS toolkit allowing for the calculation of both EI and CID spectra.^21^ Although theoretically robust, this method is hindered by the amount of time required to compute a CID mass spectra because collision events are explicitly sampled. Thus, this approach is not ideal for high throughput applications where predictions are needed in seconds, not days.

Apart from combinatorial or first-principle approaches is the application of deep learning to mass spectral prediction. The recent work of Wei et al. ^22^ predicted EI-MS spectra by training a multilayer perceptron on Extended Circular Fingerprints using a model called NEIMS. Treating EI-MS prediction as a multidimensional regression problem, they obtained a recall@1 score of 54.3%, beating CFM-EI’s score of 42.6%. To compute the recall@1 score, a predicted spectra is compared to a reference database and the score is the frequency with which the prediction’s top match is a spectrum from the target molecular ion. This led to the work of Zhu et al. who applied graph neural networks (GNNs) to higher energy collisional dissociation (HCD) MS/MS prediction.^1^ Graph neural networks bypass the use of molecular fingerprints by natively representing chemical bonding topology as input data for training and have wide ranging applications including use classifying molecules, predicting properties, and identifying molecular force fields.^23–25^ Zhu et al. showed a test set cosine similarity score of 0.51 compared to NEIMS’s cosine similarity score of 0.157 and recall@1 scores of 45.1% and 30.2% respectively.^1^ The reduction in NEIMS performance can be attributed to predicting HCD MS/MS spectra while the original implementation from Wei et al. was designed to work on EI-MS spectra.^22^

In this work, we present QC-GN^2^oMS^2^ - a GNN for MS/MS prediction using quantum chemistry (QC). As we introduce this approach, we simultaneously evaluate the performance benefit gained by adding graph edge features derived from quantum chemical methods to GNN models for high-resolution mass spectral prediction. Graph edge (bond) features were supplied from RDKit, the extended tight-binding (xTB) *ab initio* software, and the Alfabet neural network.^26–28^ We also investigate case studies in fragment assignment and explore the graph attention mechanism’s ability to explain how the GNN interprets bond breaking in MS/MS. Insights from the graph attention analysis can be used to add physics informed constraints to future models to further improve performance. Graph attention was designed to aggregate node features without reliance on rigid data structure (e.g., image convolution). ^24,29^ The equation in Figure 1b shows how to compute the attention weight *α_ij_* which determines the new vertex features **x**′_*i*_.^24^ In this graph attention method, edge features **Ea**_*ij*_ are concatenated with neighboring node features, **x**_*i*_ and **x**_*j*_. This allows edge features to participate in the gating of neighborhood information and subsequently modifies the embedded features **x**′_*i*_ output from each layer. Incorporating this attention mechanism provided minor improvement of GNN performance with the inclusion of edge features derived from quantum chemical methods.

**Figure 1:**
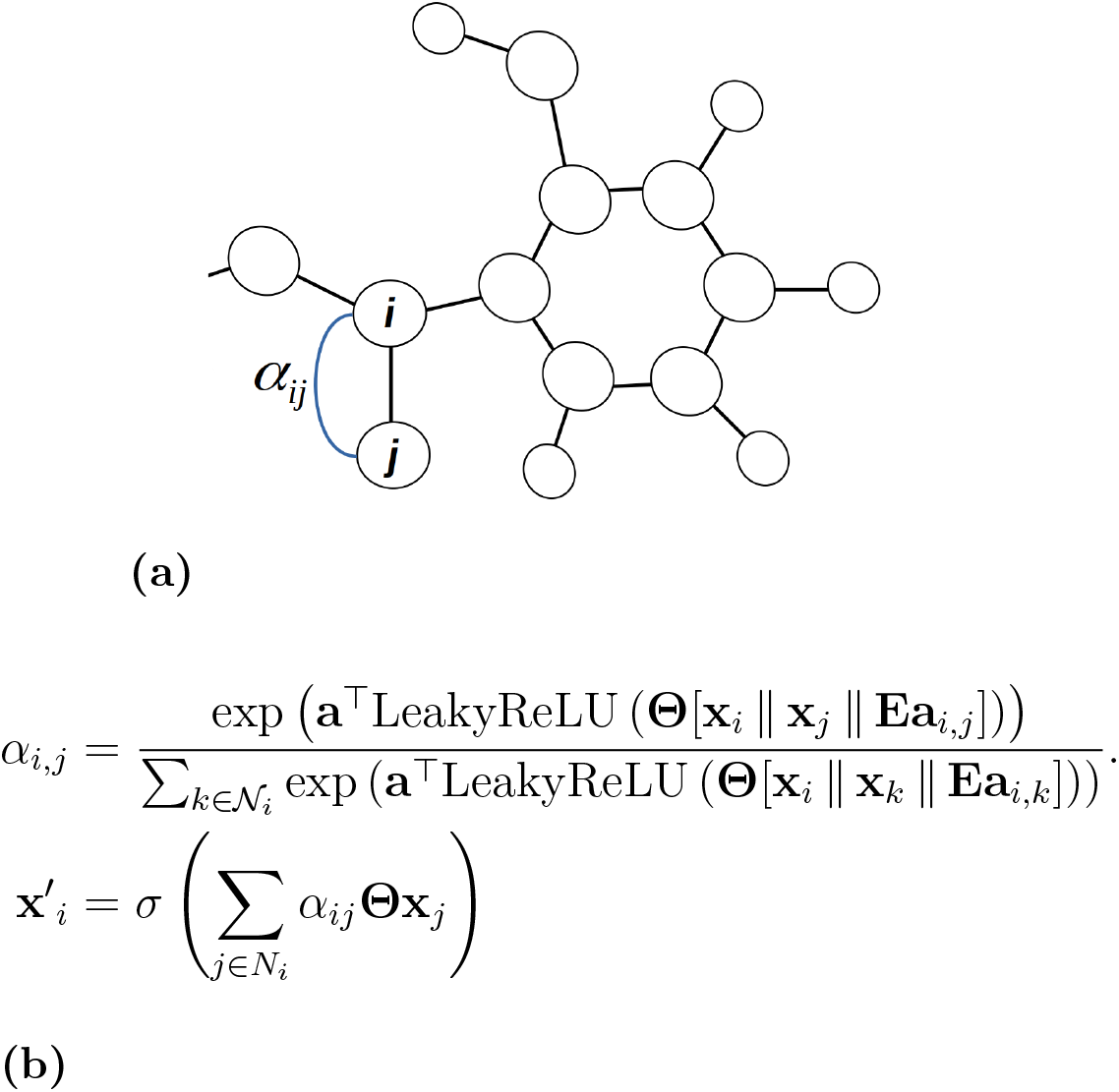
(a) Visual representation of attention coefficient computation on nodes *i* and *j*. (b) Equation for attention coefficients α_*i,j*_ where **x**_*i*_, **x**_*j*_ are node features and **Ea**_*i,j*_ are edge features. In the GATv2Conv layers, node and edge features are concatenated and multiplied by a weight matrix Θ. Each attention coefficient is normalized by summation over all nodes in neighborhood 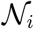 (indexed by *k*). New node features, **x**′_*i*_ are calculated from the attention weights *α_i,j_* and Θ passed through an activation function *σ*.

## Methods

Training data were aggregated from NIST20 under the following restrictions: [M+H]^+^ precursors, molecular weight *MW* ≤ 500 Da, collision energies between 30-45 eV, and instruments including the Agilent QTOF 6530 or Thermo Finnigan Elite Orbitrap. To avoid biasing during training, canonical peptides were excluded from the dataset by matching residue codes in the reference library. Because of this restriction, we do not recommend our model for MS/MS prediction on peptides; see the review by Meyer on proteomic neural networks.^30^ With these restrictions, our dataset encompasses approximately 30,000 spectra and 10,000 unique molecules. To generate the dataset used to train the GNN, spectra from NIST20 were vectorized^31^ at 50,000 dimensions to preserve high resolution data. Each floating point *m/z* value was multiplied by a constant 100 and rounded to the nearest integer index before the intensity was stored at that location in the mass spectral vector. For graph construction, InChI and SMILES strings were located in the PubChem database by cross referencing the InChIKeys in NIST20.^32^ While InChIKeys are susceptible to collisions, it is expected that mismatches are negligible over our sample size of 10,000 molecules.^33^

The GNN architecture represented in Figure 3 was built using the Torch Geometric Library.^34,35^ The primary construct in our neural network are blocks of a graph attention layer plus a linear layer. In each block, we use multi-headed attention with the assumption each head will attend to a different fragmentation pathway in the regression task. The graph attention layers in our architecture use the dynamic attention mechanism of GATv2 convolution developed in the work of Brody et al. ^24^ This mechanism has shown better performance from simple dictionary problems to the prediction of QC properties and is expected to out perform prior work using static attention for MS prediction.^1,24^ Each [GATv2Conv + Linear] layer block includes 128 channels and 8 attention heads. Blocks are separated by exponential linear unit activation functions with global mean pooling prior to the output layer. The final layer outputs a 50,000 dimension vector 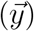 with each entry in the vector corresponding to the spectra intensity (*y_i_*) in the *m/z* bin at index i. For training, we opted for the RAdam optimizer^36^ and manually tuned the learning rate for each GNN implementation to maximize test set performance. Our loss function is a modified cosine distance function, 1 — *loss*(*x, y*), where *loss*(*x, y*) is the mean of all batch cosine similarities between predicted (*x*) and empirical (*y*) spectra. Batch size during training was 512 as this showed the best stability in training loss across models.

**Figure 2:**
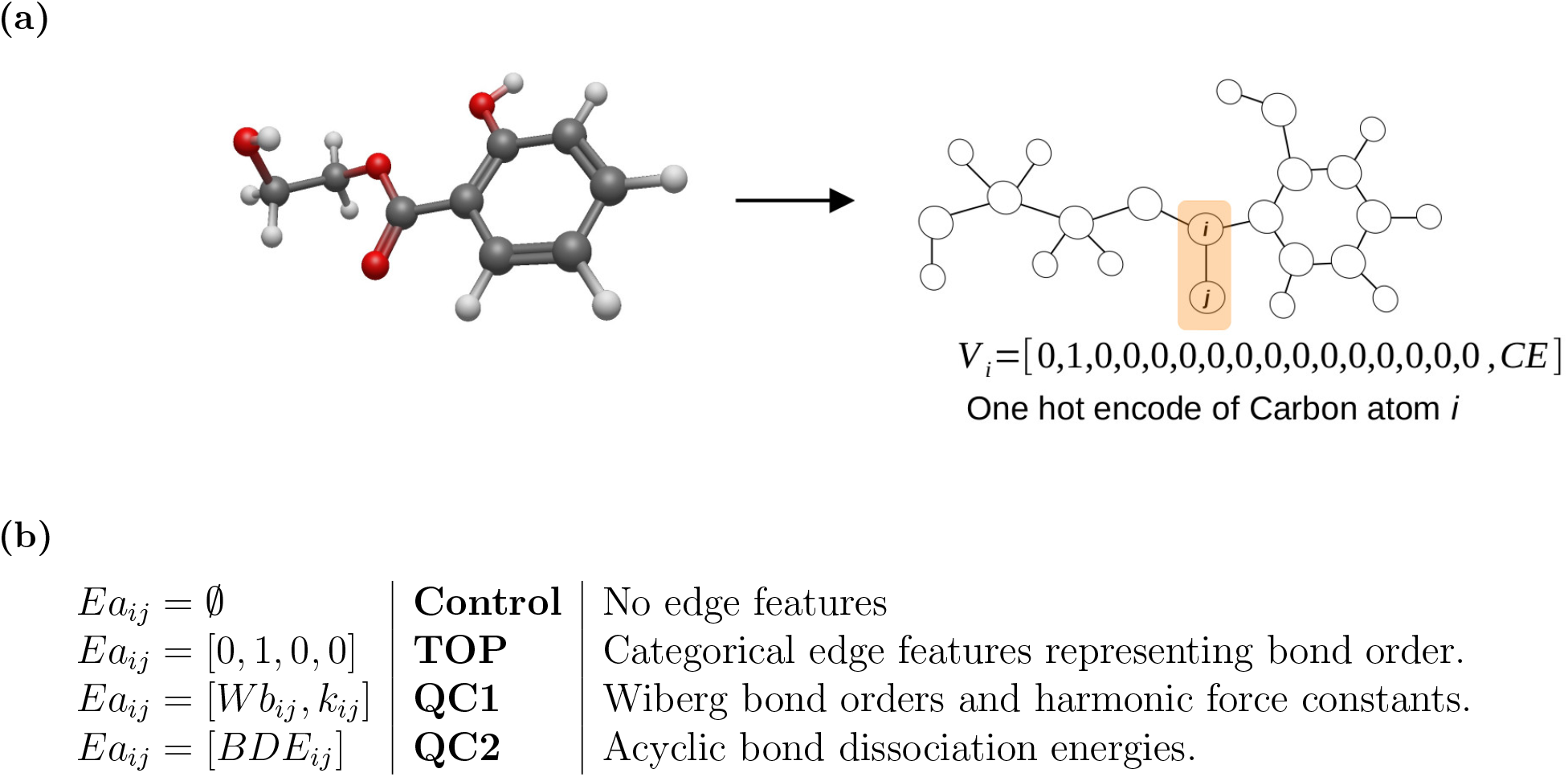
(a) Molecular graph *G* = (*V,E*) in which atoms are vertices (*V*) and bonds are edges (*E*). Hydrogens are represented explicitly in all models under consideration. Vertex features *V_i_* are a one hot encoding of the node element from the set {O, C, H, N, S, Si, P, Cl, D, F, I, Se, As, B, Br}. Node *i* is the carbonyl carbon with encoding *V_i_* = [0, 1, 0, 0, 0, 0, 0, 0, 0, 0, 0, 0, 0, 0, 0, *CE*]. Collision energy (*CE*) for each spectrum is included at the end of the vertex feature vector in electron volts (eV). (b) Listing of the models investigated in this work. Each model includes different edge features *Ea_ij_* for the graph edge *E_ij_*.

**Figure 3:**
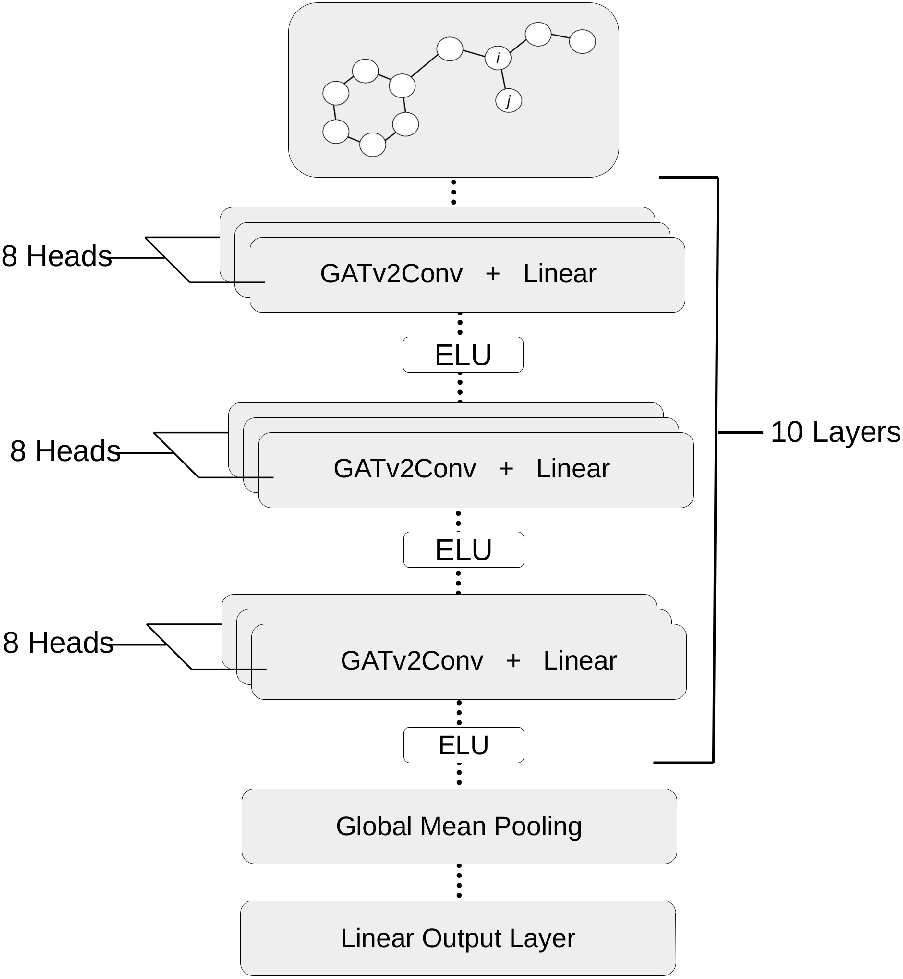
GNN architecture using multi-headed attention with 10 [GATv2Conv + Linear] blocks. Each block is separated by an exponential linear unit activation function leading to a global mean pooling and linear output layer. All layers use 128 channels. Mass spectrometry prediction is treated as a regression task with 50,000 dimensions in the output layer to preserve dataset resolution.

In the control model, molecular graphs were constructed using dataset InChI strings and RDKit.^28^ Graph vertex/atom features were encoded in a vector *V_i_* representative of element types in the set {O, C, H, N, S, Si, P, Cl, D, F, I, Se, As, B, Br}. Thus for every node there is a feature vector *V_i_* = [0, 1, 0, 0, 0,0, 0, 0, 0, 0, 0, 0, 0, 0, 0, *CE*] describing the element type (carbon) and collision energy *CE* for the graph shown in Figure 2a. Vertices are connected based on the molecule topography with the values assigned to their edge features. Unconnected vertices are assigned an edge feature 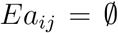 to preserves the connectivity of E without causing undesired concatenation of **x**_*i*_, **x**_*j*_, and edge features *Ea_ij_* that would affect the calculation of attention coefficients *α_ij_*. To demonstrate repeatability, the control model was trained five times with random allocations of a 1,000 molecule test set and their associated spectra. The list of molecules in each test set was stored and carried over to the three models under investigation in Figure 2b: **TOP**, **QC1**, and **QC2**. The topological **TOP** model was constructed in a similar fashion to the control representation with the addition of categorical edge features provided by RDkit bond types: single, double, triple, and aromatic.

For the **QC1** model, we attempted to improve GNN performance by providing bond force constants and Wiberg bond orders. These features were computed by using OpenBabel’s 3D conformer optimization^37^ to generate three-dimensional structures from InChI strings. Input structures were further optimized by xTB under the ‘extreme’ tolerances option that helps prevent convergence to local minima, calculation of imaginary vibrational frequencies, and non-physical Hessian values. After optimization, the coordinates, Hessian, atomic charges, Wiberg bond orders, and bond force constants were stored in a database for graph generation. Bond force constants were generated using the Seminario method. ^38^ For a bond between atoms *i* and *j* in Figure 2a, we can represent the force *δF_i_* of displacing these atoms with the partial Hessian

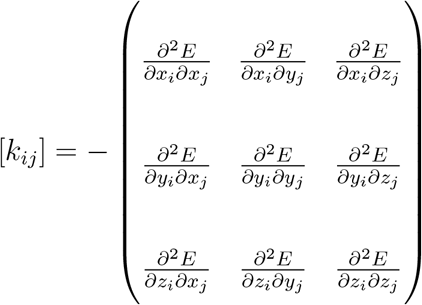

From [*k_ij_*], we compute the three eigenvalues 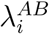 and eigenvectors 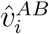 using the SciPy library.^39^ These values are used to calculate an approximation to the harmonic force constant for a particular bond by projecting each eigenvalue onto the bond vector 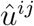:

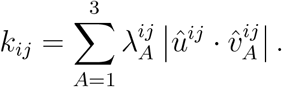

Each force constant *k_ij_* represents the stiffness of a particular bond to displacement, also thought of as how steep the basin of a harmonic potential well is for a given bond of length 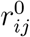,

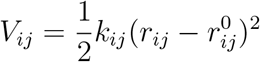

Although harmonic force constants can generalize the strength of bonds, the potential *V_ij_* can not describe the energy of bond dissociation as *r_ij_* → ∞. Thus, for an accurate description of fragmentation pathways, the transition states and bond dissociation energies of every edge in a graph would need to be known. For a single molecule, this would normally be tackled with a rigorous *ab initio* investigation of reactions using methods such as Nudged Elastic Band or Zero Temperature String.^40,41^ While descriptive of a complete fragmentation pathway, this approach becomes intractable for large datasets due to the number of fragmentation mechanisms and computational cost of saddle point search methods. As an approximation, one can ignore the contribution of transition states and only consider the thermodynamic bond dissociation enthalpy. In the work of Peter C. St. John et al., automated density functional theory calculations were performed on ~42,000 molecules to compute ~290,000 bond dissociation enthalpies.^42^ These values formed the training set for the Alfabet GNN that can predict acyclic bond dissociation energies of neutral molecules with a mean absolute error of 0.58 kcal/mol.^42^ Due to similar performance with neutral molecules, BondNet was not used for feature generation in this work.^43^ In the **QC2** representation, SMILES strings were input into the Alfabet neural network to compute edge features. Acyclic bond dissociation energies were input as floating point numbers in the edge feature matrix. Cyclic bonds, for which Alfabet does not make predictions, were input as zero in the graph edge features *Ea_ij_*.

## Results and Discussion

### Performance Metrics

Our models with different edge features (TOP, QC1, and QC2) were evaluated against a test set of molecules not seen during training. Metrics for assessing performance are shown in Figure 4 including average cosine similarity and recall@k tests. Averages shown in Figure 4a are across the five randomly allocated test sets to demonstrate stability of the GNN training protocol. The results for each model form a bimodal distribution, separating chemistries that generalize well and those that perform poorly. In the broadest terms, test distributions show poor performance with halogen functional groups, aromatic nitrogen, and oxygen in rings. In addition to the average cosine similarity score, recall@k testing was conducted with a large library to asses the quality of GNN predictions. For each recall@k test, our predicted spectra was scored against ~295,000 library spectra encompassing NIST ESI-MS/MS spectra with [M+H]^+^ precursor types and molecular weight (MW) less than or equal to 500 Daltons (Da). Scoring against a large dataset is expected to provide a more absolute metric of performance in actual reference library matching. Recall@k scores in Figure 4b indicate the probability that the correct molecule is recalled from the reference library in the first (*k* = 1) or top ten (*k* = 10) matches using GNN predicted spectra. Of the molecular representations evaluated, our best model is QC2 using bond dissociation enthalpies with metrics of 0.462, 0.102 and 0.275 for the average cosine similarity score, recall@1 and recall@10, respectively. We exclude comparisons against CFM-ID and other popular prediction tools due to differences in model design and training methodology - most popular tools were designed for nontargeted identification while our model’s loss condition rewards targeted identification. An unbiased evaluation of this model would need to include updates to fragmentation rules built as SMIRKS strings and the underlying machine-learned module.^44^ Careful examination of spectra to build novel generalized rules is outside the scope of this work.

**Figure 4:**
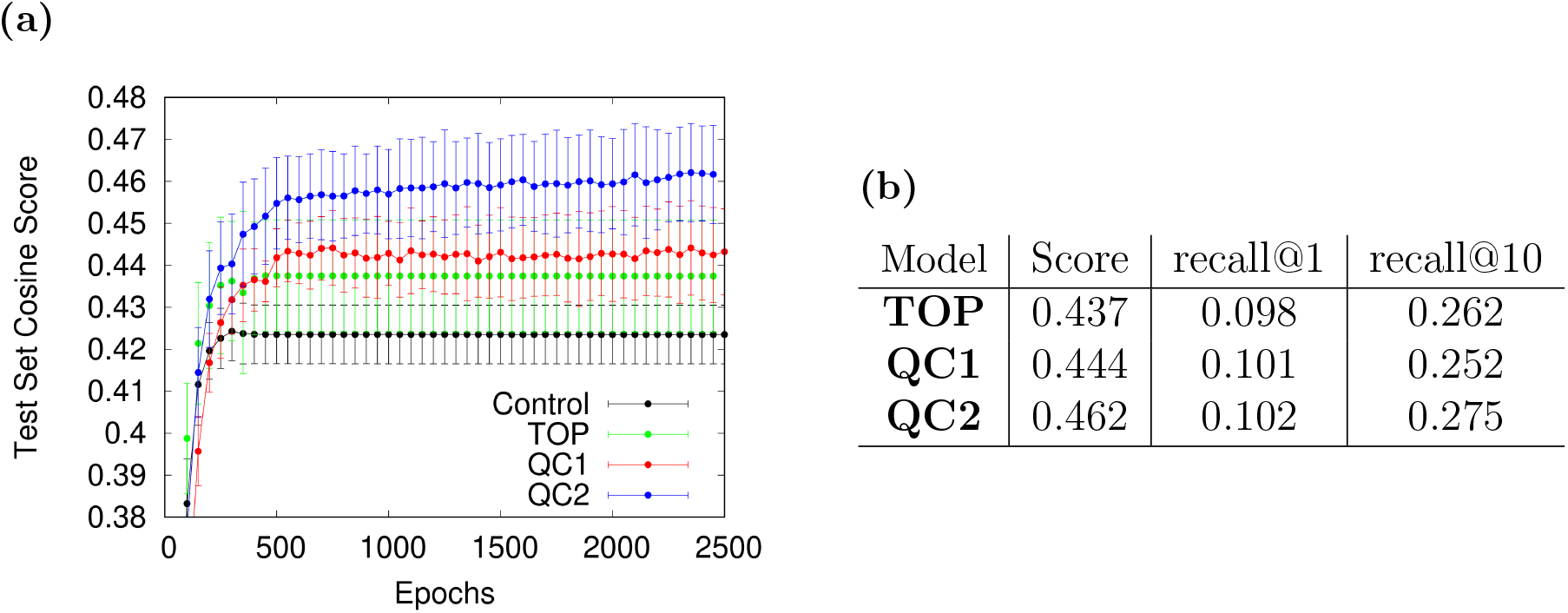
(a) Evolution of test set scores during training for Control, TOP, QC1, and QC2 models. (b) Summary of GNN performance metrics: average cosine similarity, recall@1, and recall@10 scores. Recall@k testing was conducted over 295,000 candidate spectra to represent a worst case unknown-unknown matching scenario.

Models TOP and QC1 show similar performance in test set accuracy, which could be a result of including categorical and Wiberg bond orders between the two models. During training it is possible the distribution of Wiberg bond orders is being reduced to a categorical representation during vertex feature calculation **x**′_*i*_ in each graph attention layer. In the QC2 model, we are explicitly including acyclic bond dissociation enthalpies and implicitly including neighborhood information about heterocycle content in the molecule. Although it is optimistic to consider an embedding where new node features **x**′_*i*_ are in some way representative of cyclic bond dissociation, the simplest explanation of performance gains could be due to implicit inclusion of global structure beyond nearest neighbors. Addressing this concern, we intentionally reduce bond dissociation energies to a Boolean representation of acyclic (*Ea_ij_* = 1) and cyclic (*Ea_ij_* = 0) edge character which shows an average test set cosine similarity score of 0.455, ranking this null model between the QC1 and QC2 models. Given that the null model performs worse on average than the QC2 model, we can conclude that certain quantum chemically derived edge features can make a positive impact on model performance when incorporated during training. Apart from statistical metrics, we can compare the quality of predictions across individual molecules to provide end user guidance on analyte performance.

Relative to the control representation, every addition of an edge feature made an improvement on model performance. By keeping the test sets consistent across the TOP, QC1, and QC2 models we can evaluate the per molecule performance in Figure 5. In this analysis, each point represents the cosine similarity score of a predicted spectra for a molecule in the test set at a specific collision energy. The ratio of molecules that appear above the diagonal line to molecules that appear below acts as a measure of model performance with higher numbers above the line indicating stronger performance. In the simplest case, adding estimates of bond order category (i.e., TOP) provides a performance ratio of 1.27, which indicates an improvement over pure atom connectivity. The best performance enhancement is seen with the addition of acyclic bond dissociation energies (QC2) with a ratio of 1.46. Despite this success, the majority of the improvement occurs with molecules that already have cosine similarity scores above 0.5 in the control model, which indicates that these features only work for molecules that perform well in the baseline representation. As a whole, this trend corresponds to scores shifting to above the diagonal on the top right hand corner of Figures 5b and 5c.

**Figure 5:**
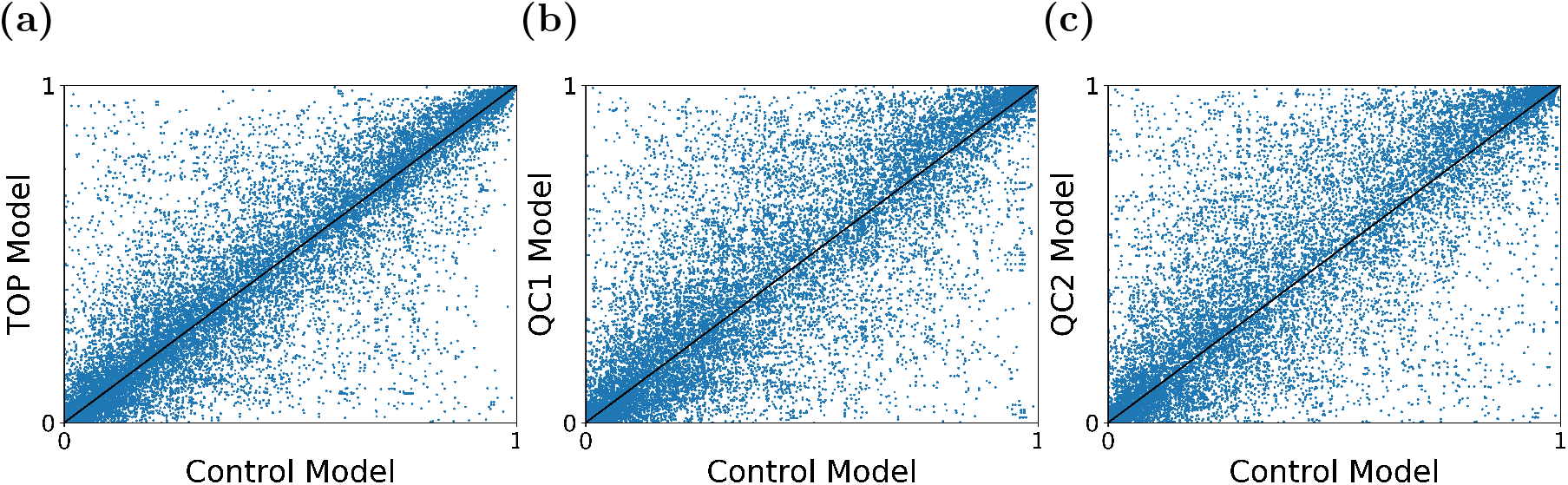
Performance of models: a) **TOP**, b) **QC1**, and c) **QC2** against control. Each point represents the cosine similarity score of an MS/MS prediction in the test set against the control representation.

Considering extreme cases, where low scoring spectra in the control model with a score of less than 0.1 switched sides to scores greater than 0.9, some rough trends emerge among models. Across representations, performance drastically improved with 2’-Deoxyguanosine, 4-(Benzhydryloxy)-1-[3-(1H-tetraazol-5-YL)propyl]piperidine, and pargyline to name a few examples. QC2 showed the best performance for these extreme cases, having 14 unique molecules dramatically shift sides in the bimodal distribution compared to 9 and 10 molecules for TOP and QC1, respectively. It should be emphasized that the above cases are anecdotal given the sample size and may not represent a systematic improvement in the GNN embedding for MS/MS prediction.

### Case Studies in GNN Fragment Embedding

Given the performance enhancement bond dissociation enthalpies provided in the QC2 model, we now turn our attention to several case studies investigating how our architecture predicts peaks for certain molecules. We isolate our investigation to predicted peaks above an intensity threshold, *IC*, and compute a node score for each identified peak, shown in figure 6. For each peak *S_i_*, we locate the rows of the linear output layer responsible for this signal and use them to form a new matrix which is multiplied by the transpose of the graph attention block output matrix. This method produces scores for each vertex summarizing vertex importance to peak *S_i_* from head *H_j_*. Our expectation is the vertices with high scores will be most representative of fragment annotation, providing a window into the GNN black box. In the test cases that follow, each vertex score is mapped to a color gradient on the molecular graph to visually represent the vertex importance distribution (VID) for peaks.

**Figure 6:**
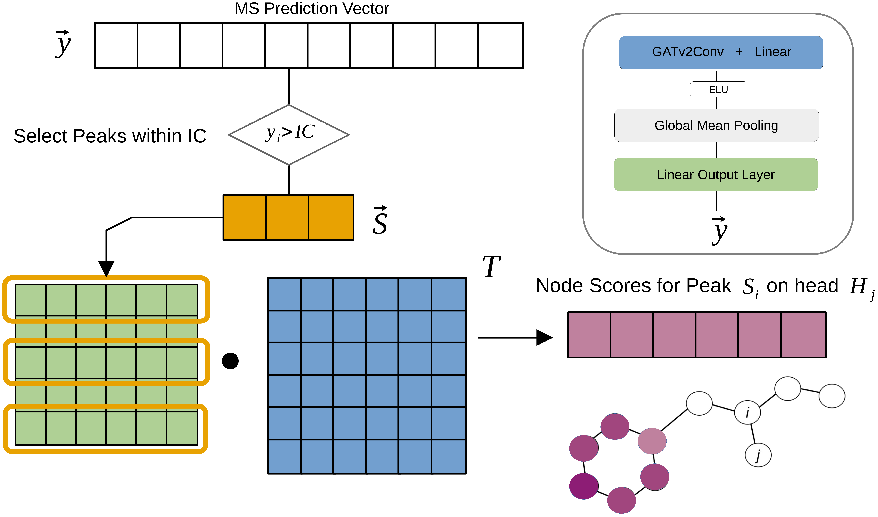
Calculation of node scores from the GNN layers to evaluate MS/MS peak to vertex assignment. Peaks are selected based on a cutoff threshold, *IC*, to limit the analysis to high intensity signals. The rows of the linear output layer (green) associated with selected peaks *S* form a new matrix which is multiplied by the transpose of the last graph attention layer output in blue. The final output of this analysis are scores for the contribution of each node at head *H_j_* to selected peak *S_i_*.

### dAMP

2’-Deoxyadenosine-5’-monophosphate (dAMP) is a fundamental metabolite and nucleotide with adenine as its base attached to a pentose ring. As a metabolite, it is produced in many pathways such as, Adenosine Deaminase Deficiency for which elevated dAMP concentrations are diagnostic.^45^ The ESI-MS/MS spectrum of dAMP collected on an Agilent QTOF 6350 at 30 eV presents with two major peaks at 136.06 and 81.03 *m/z*. The spectrum predicted by our QC2 model in Figure 7 replicates the empirical measurement well with a cosine similarity score of 0.98. Graph neural network VID of peaks S1 and S2, for each of the eight attention heads are explicitly represented in Figure 7. Across all attention heads, two categorical sub-graphs emerge, highlighting the graph vertices associated with each peak in the VID representation. In S1, greater weight is given to the pentose ring with some partial importance placed on the adenine base. Peak S2 shows a greater importance on the adenine base partially cleaving the pentose ring during vertex assignment and losing an amino group in five of the eight attention heads. Manually accounting for weights of dAMP functional groups, the VID across peaks appears to align well with the molecular weight of the adenine base 150.13 g/mol less an amino group accounting for peak S2 and partial segments of the pentose ring (S1). The quality of this alignment is likely due to the bimodal VID of scores separating these functional groups.

**Figure 7:**
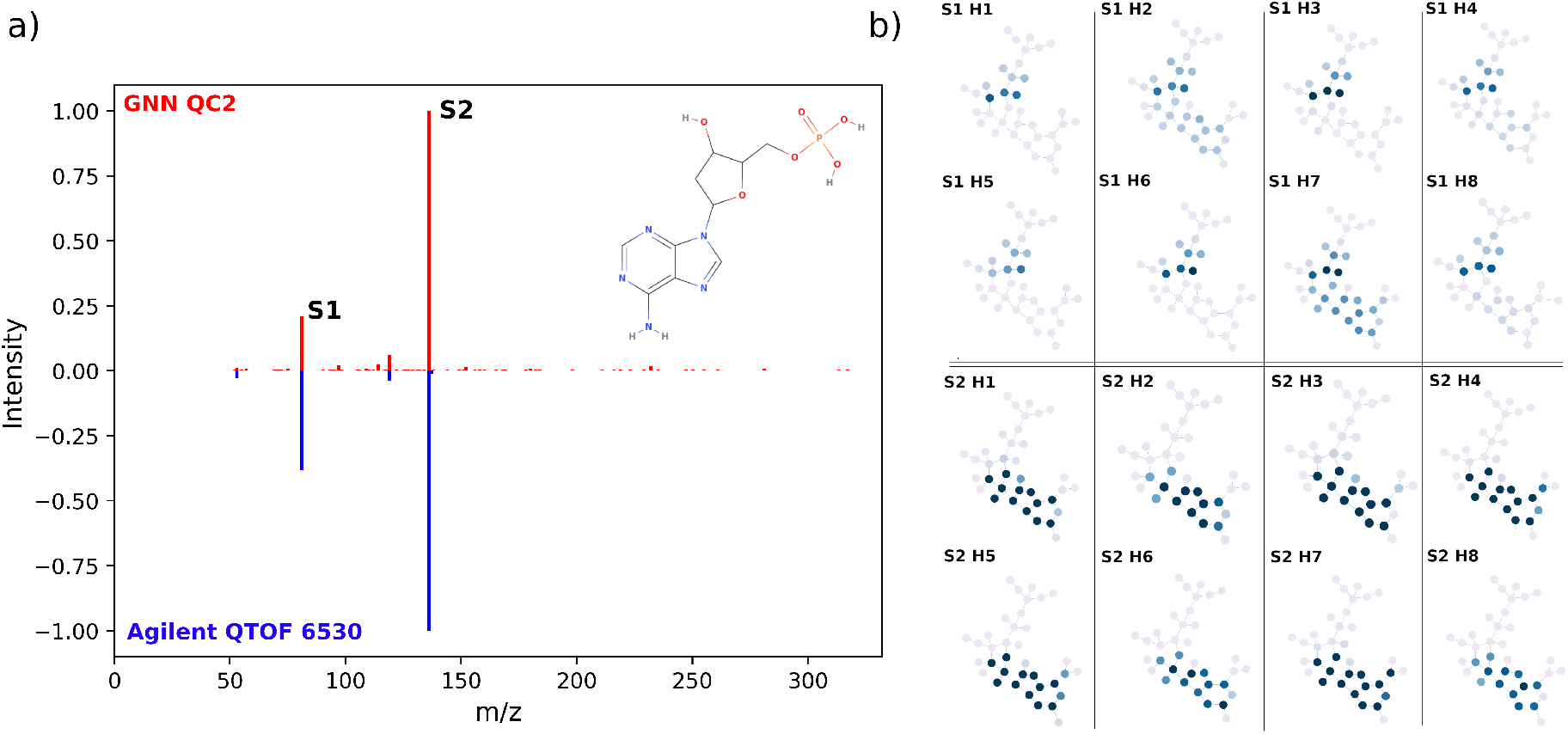
2’-Deoxyadenosine-5’-monophosphate: (a) MS/MS spectrum predicted with the QC2 model at 30 eV and a cosine similarity score of 0.98. (b) VIDs indicating the importance of an atom in the *Sn* peak. Scores on each *Hm* head are shown for selected peaks *S*1 (81.03 *m/z*) and *S*2 (136.06 *m/z*). VIDs for peaks S1 and S2 show strong correlation with segments of the pentose ring and adenine base. *Sn Hm* indicates the *n^th^* peak and *m^th^* attention head.

### 2-Chloro-L-phenylalanine

2-Chloro-L-phenylalanine is a non canonical amino acid often used as an internal standard in metabolic studies. The predicted spectrum is shown in Figure 8a with the corresponding empirical spectra plotted below in blue. Performance against the empirical spectra is quite poor with a cosine similarity score of 0.016 and can likely be attributed to the addition of a chlorine substituent on the aromatic ring. Vertex scores in figure 8b shows a reliance on breaking of the aromatic ring for base peak S1, generating an overestimate at 125.02 *m/z*. For this VID, it appears the GNN is biased by a under representation of halogenated substituents relative to phenyl functional groups in the training data. An *in silico* investigation of 2-Chloro-L-phenylalanine CID fragmentation pathways with QCxMS shows a general consensus in the assignment of the 118.06 *m/z* fragment. Generation of this fragment generally involves breaking first the C_*α*_-C_*O*_ bond followed by removal of the aromatic halogen. Given this pathway, another explanation of poor performance is the inability of the neural network to handle multi-step fragmentation pathways especially in cases where chemical neighborhoods are well preserved in other spectra (i.e., phenyl at 77 *m/z*). Despite the representation of empirically consistent fragments in the 45 eV QCxMS spectra, overall performance of this method is also quite poor with a cosine similarity score of 0.0452 relative to the empirical spectra. Head to tail plots of this *ab initio* spectrum are available in the supplemental information and displays an over abundance of high and low m/z peaks relative to the empirical base peak. The inaccuracy of this first principles method can be due to a number of factors including the level of theory, precursor protonation state selection, or calibration of the collision energy settings for a particular system. Given the under sampling of empirical fragmentation pathways, parameter sensitivity, and computational resources we did not pursue a rigorous performance evaluation of QCxMS against our GNN models.

**Figure 8:**
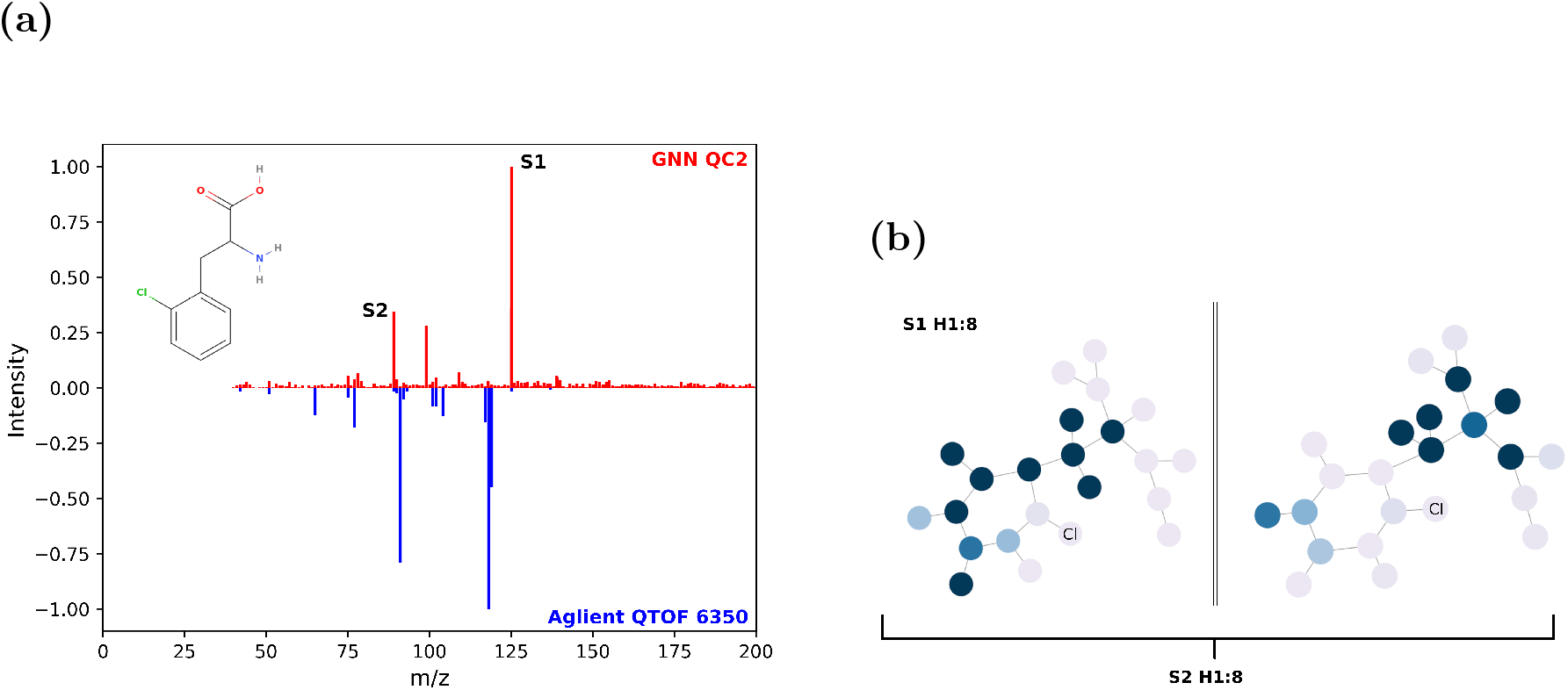
2-Chloro-L-phenylalanine, a case of poor performance. ESI-MS/MS spectrum predicted at 45 eV with the QC2 model and a cosine score of 0.016. (a) Predicted (red) and empirical spectrum (blue) collected on a Aglient QTOF 6350. (b) VIDs consolidated across heads 1-8 for peaks S1 (125.02 *m/z*) and S2 (89.04 *m/z*).

### Hydrocortisone

Hydrocortisone is a medicinal application of the hormone cortisol used to treat many diseases, the most common being dermatitis thorough topical application. Containing multiple ring structures and chiral centers, the MS/MS spectrum is especially rich in the number of fragment peaks. Given the number of coupled fragmentation pathways, this spectrum would be difficult to approximate in *silico*. The base peak for this spectra at 121.06 *m/z* is assigned to the S2 embedding in Figure 9b. This is in disagreement with the NIST MS Interpreter annotation assigning this fragment to the cyclopentane heterocycle. Looking to CFM-ID annotation, the MS/MS embeddings S1/S2 show better agreement with a focus on the cyclohexenone/cyclohexane rings. Aside from peak annotation, it could be concluded the graph attention layers are able to mix neighborhood representations to better approximate multiple fragmentation pathways in complex spectra, hence the disjointed VID in Figure 9b.

**Figure 9:**
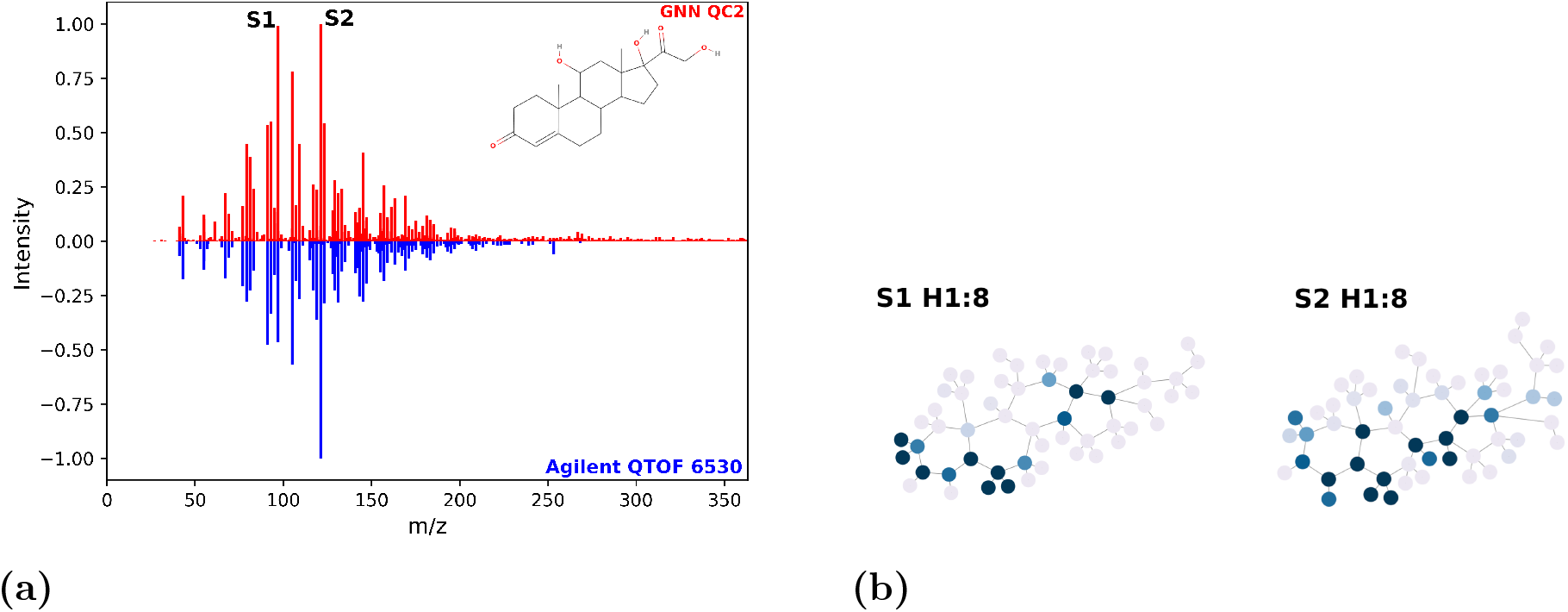
(a) Predicted ESI-MS/MS spectrum of hydrocortisone at 45 eV using the QC2 model with a cosine score of 0.88 relative to the empirical spectra. (b) Summarized VIDs of peaks S1 (97.06 *m/z*) and S2 (121.06 *m/z*).

## Conclusion

Graph neural networks have shown great promise in molecule classification, property regression, and mass spectral prediction in part due to their ability to represent chemical structure. In this work, we presented three QC-GN^2^oMS^2^ models that vary the edge features of the input graphs including bond order (**TOP**), Wiberg bond orders with harmonic force constants (**QC1**), and bond dissociation enthalpies (**QC2**). Out of the representations considered, the QC2 model showed the best performance in terms of average cosine similarity and recall@k scores. Exploration of the GNN embedding showed chemically reasonable fragment annotation for simple and complex spectra. In our example of 2’-Deoxyadenosine-5’-monophosphate, the predicted spectrum shows good agreement with the NIST database and fragment annotation pointing to N-C glycosidic bond cleavage commonly observed in dissociation of protonated nucleosides.^46^ The bimodal distribution of test set scores illustrates that some chemical topologies perform poorly such as halogens, nitro groups, aromatic nitrogen or cage structures. A potential rational for poor performance of certain functional groups are under representation in the training data or the charge dependent shifts in molecular geometries, that affect fragmentation pathways, force constants or bond dissociation energies.^43^ Despite difficulties with certain functional groups we have demonstrated the net benefit quantum chemical features provide graphical neural nets by including BDEs as edge features. Future work includes architecture development for improved mechanistic attention, combining QC1 and QC2 features, and the combinatorial breaking of heterocycles to compute average cyclic bond dissociation enthalpies with Alfabet.

## Acknowledgement

This work was supported by the PNNL Laboratory Directed Research and Development Program and is a contribution of the *m/q* Initiative. PNNL is a multi-program national laboratory operated by Battelle for the DOE under Contract DE-AC05-76RLO 1830.

## TOC Graphic

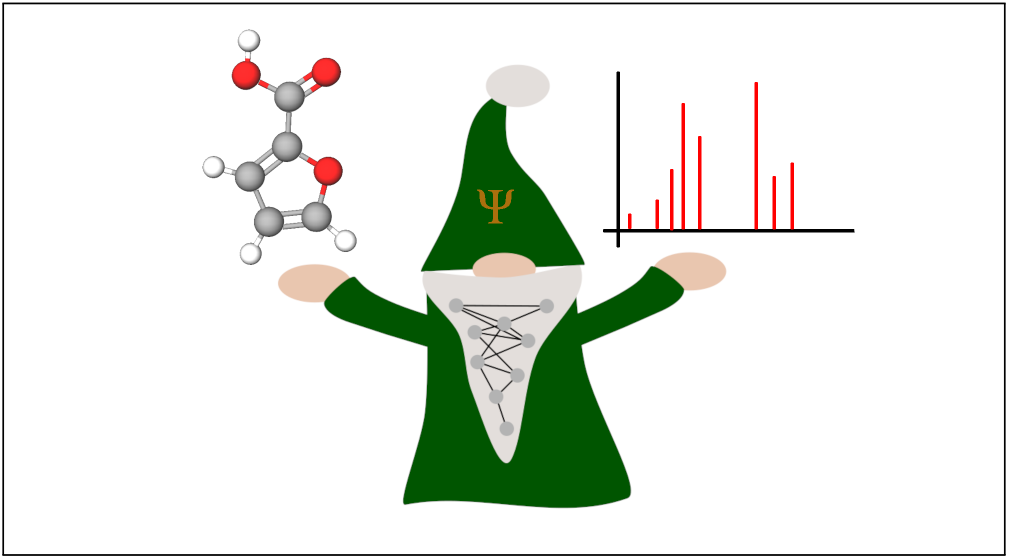

