## Supplemental Figures for "QC-GN^2^oMS^2^: a Graph Neural Net for High Resolution Mass Spectra Prediction"

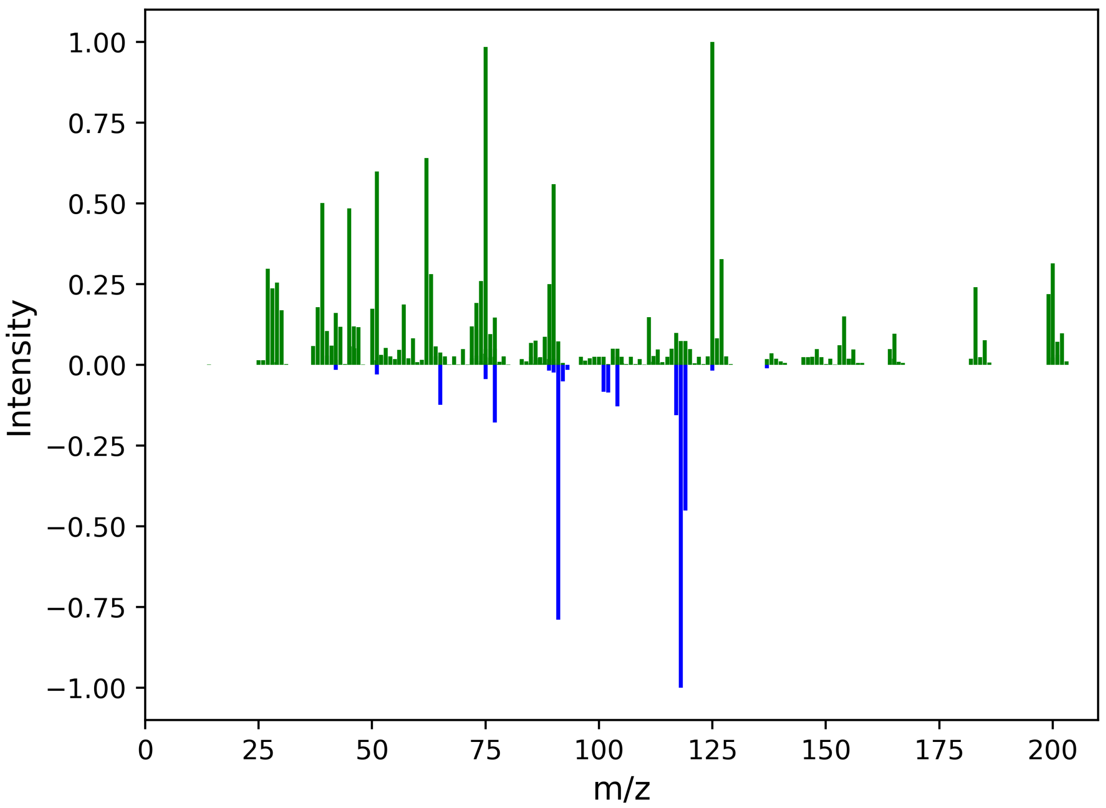


Figure 1: 2-Chloro-L-phenylalanine QCxMS Spectra calculated at 45eV compared against the ESI-MS/MS [M+H]+ spectra collected on a Agilent QTOF 6530
